# Two phases of aging separated by the Smurf transition as a public path to death

**DOI:** 10.1101/028183

**Authors:** E. Dambroise, L. Monnier, L. Ruisheng, H. Aguilaniu, J-S Joly, H. Tricoire, M. Rera

**Affiliations:** Unité de Biologie Fonctionnelle et Adaptative (BFA) UMR8251 - CNRS - Université Diderot, Sorbonne Paris Cité, 4 Rue Marie Andrée Lagroua Weill Hallé, 75013, Paris, France; Equipe CASBAH “Comparative Analysis of Stem cells, Brain Anatomy and Homeostasis”, Neuroscience Paris-Saclay Institute (Neuro-PSI), UMR 9197, CNRS, Gif-sur-Yvette, France; Ecole Normale Supérieure de Lyon - CNRS - Université de Lyon Claude Bernard - Institut de Génomique Fonctionnelle de Lyon /UMR5262 46, Allée d’Italie, 69364 Lyon, France

## Abstract

Aging’s most obvious characteristic is the time dependent increase of an individual’s probability to die. This lifelong process is accompanied by a large number of molecular and physiological changes. Although numerous genes involved in aging have been identified in the past decades its leading factors have yet to be determined. To identify the very processes driving aging we have developed in the past years an assay to identify physiologically old individuals in a synchronized population of *Drosophila melanogaster*. Those individuals show an age-dependent increase of intestinal permeability followed by a high risk of death. Here we show that this physiological marker of aging is conserved in 3 invertebrate species *Drosophila mojavensis, Drosophila virilis, Caenorhabditis elegans* and 1 vertebrate specie *Danio rerio.* Our findings suggest that intestinal barrier dysfunction may be an important event in the aging process conserved across a broad range of species, thus raising the possibility that it may also be the case in *Homo sapiens*.

## Introduction

Based on the evolutionary links existing between species, it is common in biology to address mechanistic questions using model organisms. Organisms such as *Drosophila melanogaster* or *Caenorhabditis elegans* have been extensively used for deciphering genetic and molecular pathways in many biological processes, including aging. In fact, the first ‘longevity gene’ has been identified in *C. elegans*^1^, which led to the discovery of the insulin pathway as a major conserved regulator of aging in many species. Since then, many more genes were shown as playing a role in determining longevity^2-6^.

Nevertheless, numerous genomic rearrangements and organism specificities that occurred during evolution might lead to the identification of mechanisms that cannot be transposed from one organism to another. For example, the complexification of insulin signaling pathway in mammals compared to ecdysozoans’ might render pro-longevity genetic interventions deleterious or neutral when transposing longevity treatments identified in the latter organisms^7^.

Thus, one critical step towards identifying the leading factors driving aging would be to switch from an all-genetic paradigm to a more physiological paradigm. Such a switch would be possible through identifying an evolutionarily conserved sequence of physiological and/or molecular events occurring during normal aging – in contrast to pathological aging. We interpret the recently described Smurf phenotype^3^ as such a sequence in drosophila. The Smurf phenotype is a dramatic increase of intestinal permeability that can be observed *in vivo* using a non-invasive assay. The proportion of individuals showing that phenotype increases quasi linearly as a function of chronological age and every individual will undergo this change prior to death from non-pathological causes, in other words ‘occurring during normal aging’. Individuals exhibiting this phenotype show a dramatic increase of their probability to die compared to their non-Smurf siblings whatever their chronological age. Finally, this phenotype is accompanied by strong modifications of numerous other molecular and physiological phenotypes that are commonly described as being hallmarks of aging (i.e. reduced locomotion, reduced energy stores, increased inflammation,…)^8^. This complex transition occurs in every fly during normal aging in a manner that seems extremely stereotyped. We recently provided a theoretical framework to these aging related features in a new model of aging (2PAC) that describes aging in individuals as being composed of two successive phases each associated with distinct properties and separated by a dramatic physiological transition^9^.

We hypothesized that the Smurf phenotype could be conserved across evolutionarily distant organisms as an easily detectable harbinger of death occurring in physiologically old individuals. To test our hypothesis, we tested the validity of the following three assumptions on three model organisms currently used in the study of aging processes: i) in a given synchronized population some individuals show increased intestinal permeability characterized by the Smurf phenotype, ii) the proportion of Smurfs in the population is age-related and ultimately iii) the presence of the Smurf phenotype is associated to a high risk of death. We validated these assumptions in all the studied organisms, including a vertebrate species. Thus, we propose that the Smurf phenotype is an evolutionarily conserved marker of advanced physiological age. Further identification and molecular characterization of this aging related phase transition in distant species – including *Homo sapiens –* as well as in longevity modifying experiments would shed new lights on the very processes driving aging.

## Results

It was reported by Rera et al. in 2011 that a dramatic increase of intestinal permeability occurs in *Drosophila melanogaster* during aging in normal condition. The assay presented in this article uses a blue food dye to detect increased intestinal permeability *in vivo*. A blue coloration throughout the body marked the positive individuals, which were referred to as ‘Smurfs’ from then on. Interestingly, the authors showed that genetic or physiological interventions increasing lifespan in flies significantly decreases the proportion of Smurfs compared to the control population at any given chronological age^3^. This apparent link between the age-related increase of intestinal permeability and lifespan led them to more thoroughly analyze the Smurf phenotype^8^. This phenotype allows the identification of individuals that are about to die of natural death amongst a population of synchronized *Drosophila melanogaster* individuals and those individuals show numerous other hallmarks of aging. Such a stereotyped way to die is unexpected; this could indicate a physiological phenomenon crucial during normal aging. Here we propose to test the hypothesis that such an important phenomenon should be evolutionarily conserved.

We chose to search for such a ‘Smurf transition’ in two other drosophila species, *Drosophila mojavensis* and *Drosophila virilis* whose last common ancestor with *Drosophila melanogaster* existed approximately 50 million years ago, the nematode *Caenorhabditis elegans* whose divergence time with *D. melanogaster* is around 750 million years, and finally the vertebrate zebrafish *Danio rerio,* which diverged from *D. melanogaster* around 850 million years ago. Divergence times were obtained on timetree.org. Besides the broad time scales separating those species from *Drosophila melanogaster*, they are also characterized by a broad range of life expectancies at birth – from 10 days to 40 months (Fig. 1).

**Figure 1:**
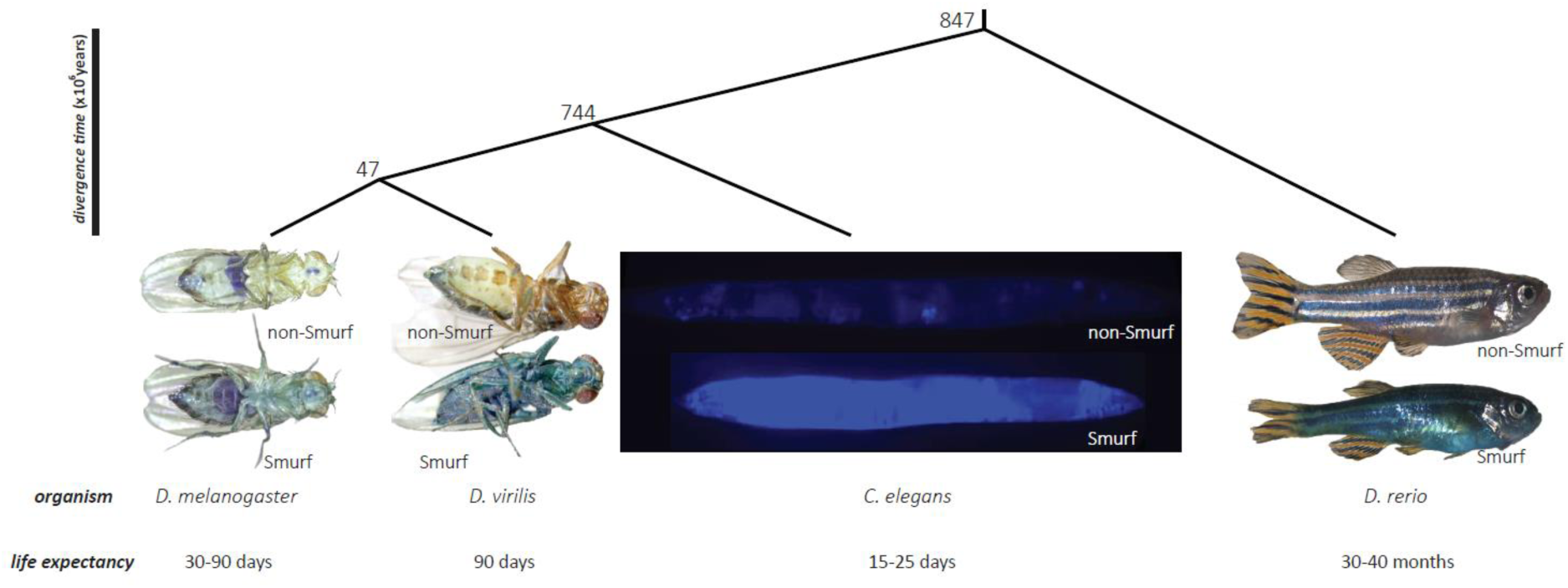
The intestinal permeability detected using the Smurf assay is evolutionarily conserved. (A) Phylogenic tree of the three studied organism. (B) We previously described the Smurf phenotype as the dramatic age-related increase of intestinal permeability in *Drosophila melanogaster* observed by feeding individuals a non-absorbed food dye. We show here that it is possible to observe a similar intestinal permeability in evolutionarily distant organisms, two other drosophila species *D. virilis* and *D. mojavensis* as well as the nematode *Caenorhabditis elegans* and the vertebrate *Danio rerio.* The pictures presented are of female drosophila, hermaphrodite nematodes and female zebrafish.

### Smurfs do exist in drosophila, nematodes and zebrafish

We investigated whether Smurf-like animals could be observed in individuals from populations of these evolutionarily distant organisms. We first tested the possibility to observe the Smurf phenotype by searching for it in old individuals of each species, respectively 40 days in flies, 12 days in nematodes and 48 months in the zebrafish. We adapted the Smurf Assay (SA) protocol to the specificities of each organism (see *material and methods*). Notably, to facilitate detection in nematodes, the blue dye #1 was replaced by fluorescein that was previously reported as allowing the detection of Smurfs in *D. melanogaster*^8^. For each tested species, we could identify individuals showing extended dye coloration throughout their bodies. Moreover, we observed an heterogeneity in a given population, with only a fraction of the individuals exhibiting increased dye level outside the intestine. Thus, at least in old animals, it is possible to identify, using SA, individuals with increased intestinal permeability in *Drosophila melanogaster*, *Drosophila virilis*, *Drosophila mojavensis*, *Caenorhabditis elegans* and the vertebrate *Danio rerio* (Fig. 1).

### Smurfs proportion increases linearly as a function of chronological age

An important result of our previous work in *D. melanogaster*, is that the proportion of Smurf individuals in a synchronized population is growing linearly as a function of chronological time **t**: for t > t_0_, the proportion S of Smurf individuals in the population varies as S = SIR^*^(t-t_0_) where the constant SIR is called the Smurf Increase Rate and **t_0_** is the first age at which one can detect Smurfs in the population ^9^. Two populations exhibiting similar longevity curves have non-significantly different **SIRs** and **t_0_**. On the other hand, populations showing significantly different lifespans can either have different **SIRs**^8^ or **t_0_**^9^.

We first searched for such an age-dependent linear increase of the Smurf proportion in synchronized populations of *D. virilis* and *D. mojavensis* both characterized by significantly different lifespans (Fig. 2A) by testing them using the SA every 10 days as previously described in *D. melanogaster*^8^. First, we could observe that for each species, the proportion of Smurfs increases as a function of time, in a quasi linear manner (Fig. 2B). Moreover, as previously described in *D. melanogaster*, the population showing the shortest lifespan has the highest SIR. Not only the *D. mojavensis* population has the shortest life expectancy (T_50_ = 42 days) but it also starts showing dead individuals earlier (21 days) than the longer lived *D. virilis* population (38 days).

**Figure 2:**
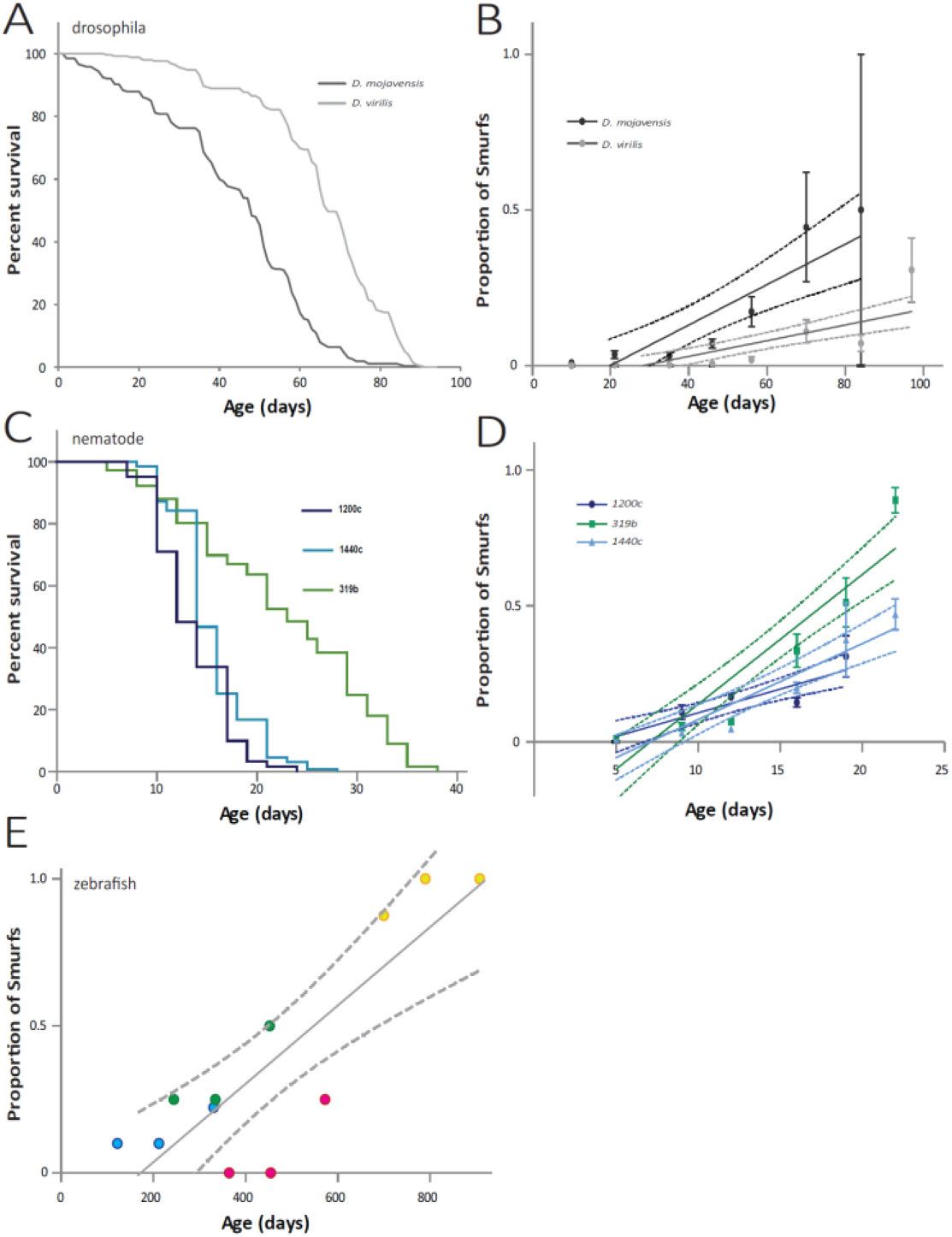
Proportion of individuals showing the Smurf phenotype increases in an age-dependent manner as previously described in *D. melanogaster*. (A) The lifespan of *D. mojavensis* females (T_50_ = 47.7 days, N = 265) is significantly reduced compared to that of *D. virilis* females (T_50_ = 66.8 days, N = 252) (log rank p < 0.0001). (B) The Smurf Increase Rates (SIRs) of both populations are significantly non-null (p_Ftest_ < 0.0001). The SIR of the longest lived population of *D. virilis* (SIR*_virilis_* = 0.002516 ± 0.0004535, R^²^ = 0.2570) is significantly lower (p_Ftest_ = 0.002892) than the SIR of the shorter lived *D. mojavensis* population (SIR*_mojavensis_* = 0.002516 ± 0.0004535, R^²^ = 0.2938). This negative correlation between SIR and life expectancy was previously showed in *D. melanogaster*. (C) The lifespan of three populations of *C. elegans* hermaphrodites was assessed, 319b (T_50_ = 19 days, N = 180), 1440c (T_50_ = 14 days, N = 131) and 1200c (T_50_ = 12 days, N = 134). (D) The SIR of 1440c (SIR_1440c_ = 0.02791 ± 0.003904, R^²^ = 0.7290) is not significantly different (p_Ftest_ = 0.6212) from the SIR_1200c_ (0.01048 ± 0.02440, R^²^ = 0.6383) as expected from their similar lifespan curves. The SIR_319b_ (0.04770 ± 0.005246, R^²^ = 0.8212) is significantly different from SIR1440c (p_Ftest_ = 0.004342). Because the slopes differ so much, it is not possible to test whether the y-axis intercepts differ significantly. (E) In the zebrafish males too, the SIR (0.001326 ± 0.0002557, R^²^ = 0.7291) is significantly non-null (p_Ftest_ = 0.0004). The four independent groups used for the Smurf Assay are represented by four different colours (blue, green, pink and orange). The number of values for calculating each SIR is above 12. Error bars represent s.e.m. Dashed lines represent I.C.95.

Then we examined whether an age-dependent linear increase of the proportion of individuals characterized by an increased intestinal permeability can be observed in nematodes. First and importantly we did not observe any deleterious effects of fluorescein on worm survival when we compared two N2 populations, one reared on standard food the other on standard food complemented with fluorescein (sup Fig. 1A). Then we assayed for Smurfs three natural populations of nematodes characterized by similar lifespan (1200c, T_50_ = 12 days and 1440c, T_50_ = 14 days) or significantly different ones (319b, T_50_ = 19 days)^10^ (Fig. 2C). All three lines showed an age-dependent linear increase of individuals showing an extended dye labeling of the whole body post SA. Moreover, as previously described for the *D. melanogaster* lines *w*^1118^ and CantonS characterized by overlapping longevity curves^8^, the *C. elegans* lines 1200c and 1440c show SIRs and t_0_ that are not significantly different with respectively, p_slopes_ = 0.06212 and p_t0_ = 0.9928. On the other hand, the long-lived 319b line shows a SIR that is significantly higher than SIR_1200c_ and SIR_1440c_. However, this higher Smurf Increase Rate starts three days after the first Smurfs appeared in the shortest lived lines (Fig. 2D). But interestingly, the latter population shows the first Smurf individuals much later as expected from the 2PAC model^9^.

Finally, we decided to test the age-dependent increase of the probability to observe intestinal permeability in zebrafish. To overcome the major problem of this model which is its relatively long life expectancy, we studied the proportion of Smurfs in four groups of both males and females *D. rerio* (AB genotype) of different ages (respectively 4, 8, 12 and 23 months) that were then assayed again 3 and 7 months later. As previously described in drosophila and nematode, the proportion of Smurfs increases linearly in male fishes as a function of time (p = 0.0004, Fig. 2E). In females, this correlation was not significant (p = 0.0597, sup Fig. 1B) although a trend is visible. This may be due to the low number of individuals in the different categories.

### Smurfness as a ‘harbinger of death’ is evolutionarily conserved

One of the most striking characteristic of Smurf individuals previously described in drosophila is the high risk of impending death they exhibit compared to their age-matched counterparts in a given population^8^. So we decided to verify whether Smurf individuals were also committed to die in the other organisms we studied in this article. As described in ^9^, to maximize the number of individuals and obtain a representative picture of a Smurf’s lifespan, we grouped all the Smurfs obtained throughout life for each organism to plot each Smurf survival curve.

The survival curves of Smurfs from *D. virilis* and *D. mojavensis* are highly similar with both T_50_s equal to 2.5 days (Fig. 3A) although the life expectancy at birth is significantly different in those two species (Fig. 2A). Interestingly, this T_50_ value is close to what was previously described for *D. melanogaster* females in ^8^ and is identical to what we recently described using a different SA protocol^9^. For nematodes, we followed a similar protocol and could find that Smurf survival curves are completely overlapping for the two strains, 1440c (T_50_ = 4.2 days) and 319b (T_50_ = 4.1 Days) (Fig. 3B) although their life expectancies at birth are significantly different (Fig. 2C). Finally, although no mortality could be observed, so far, in neither Smurfs nor non-Smurfs from the three youngest groups of zebrafish, more than half of Smurfs from the oldest group died after 8 months (4 out of 7 individuals) while non-Smurfs from the same age didn’t show any mortality (n = 9) (Fig. 3C). In regard to this data, a statistically non-significant trend of decreasing life expectancies of Smurf flies of increasing chronological age has been observed in Smurfs flies of different ages^8,9^ although the overall remaining lifespan of such individuals is pretty standard. It is possible to imagine that the few days of difference observed in the T_50_ of Smurfs from different chronological ages observed in drosophila or nematodes become weeks or even months in a long lived species.

**Figure 3:**
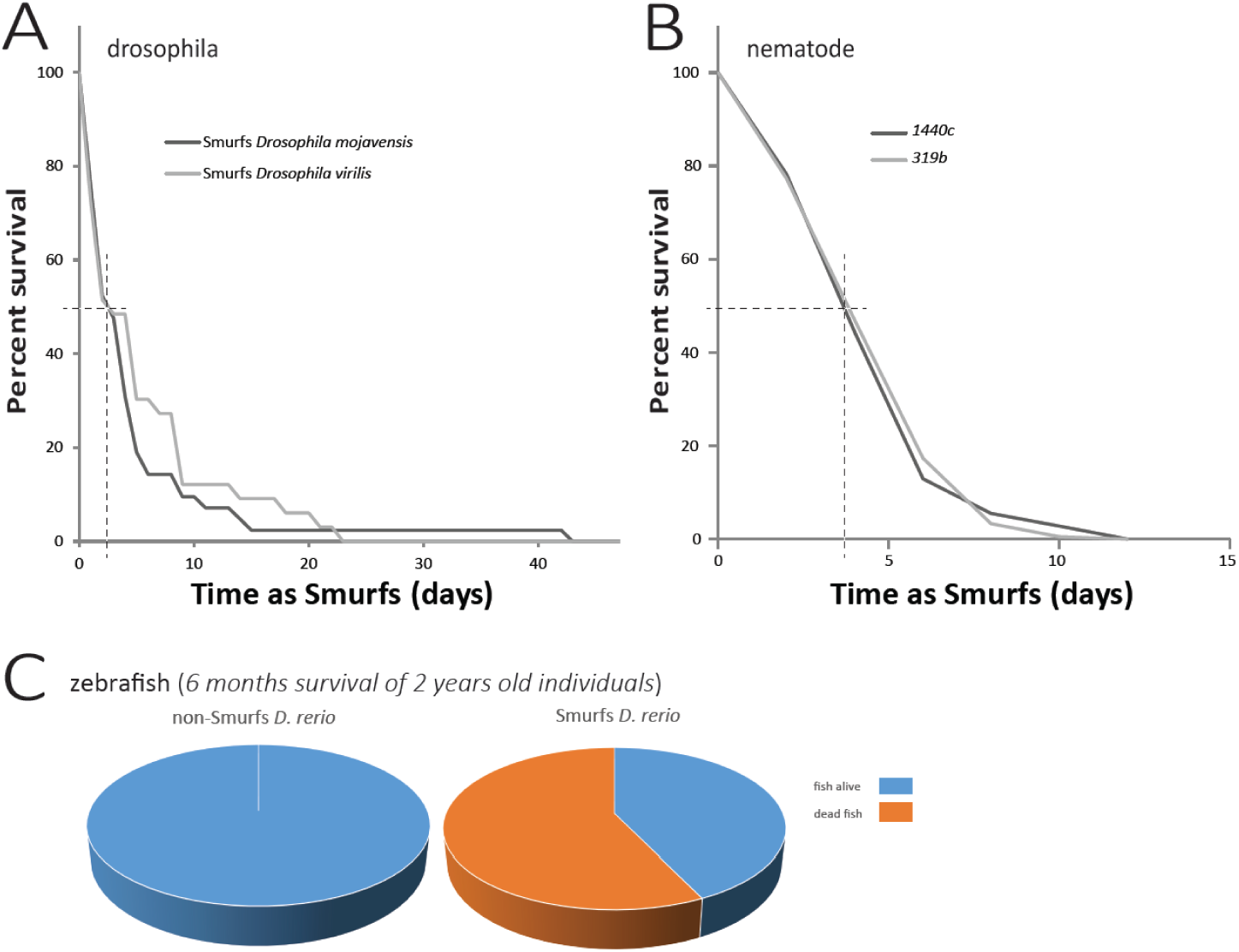
The Smurf phenotype is an evolutionarily conserved harbinger of death occurring during normal aging. (A) *D. mojavensis* females Smurfs remaining lifespan (T_50_ = 2.5 days, N = 45) is not significantly different to that of *D. virilis* females Smurfs (T_50_ = 2.5 days, N = 33) although both species show significantly different life expectancies at birth. (B) Smurf from *C. elegans* lines 1440c (T_50_ = 4.2 days, N = 45) and 319b (T_50_ = 4.1 days, N = 45) show identical survival curves (p_logrank_ = 0.671) although both lines are characterized by significantly different life expectancies at birth. (C) Smurf zebrafish males that were scored at 2.5 years of age (N = 7) show a significantly higher risk of death 6 months after the scoring (N = 4) than their age-matched non-Smurf siblings (0/9) (p_binomial_ = p < 2.2e-16).

## Discussion

A major challenge in the aging field is to identify the chain of events responsible for an organism to become unable to cope with environmental or internal cues that are usually managed well. In a recent article^8^, Rera and colleagues identified a physiological defect occurring in the aging drosophila that could be directly related to a high risk of impending death from ‘natural causes’. We interpret the Smurf phenotype described in this paper as one manifestation of such a sequence of events. As it allows the identification of individuals that are about to die from natural causes, we believe that this phenotype may allow a massive change of paradigm in how to study aging. Indeed, the strategy that has been privileged since the identification of the first gene able to modulate longevity is to search for genetic or pharmacological interventions able to increase healthspan. If the Smurf phenotype were to be evolutionarily conserved, it would allow focusing on the only element common to all aging organism, their death from ‘natural causes’.

In the present article, we decided to search for a similar phenotype in evolutionarily distant organisms. First, in two other drosophila species, *D. mojavensis* and *D. virilis*, then in the ecdysozoan *Caenorhabditis elegans* and the vertebrate *Danio rerio*, we show that it is possible to detect an increased intestinal permeability to a dye *in vivo* in species that have diverged from 900 to 50 million years ago. Secondly, we showed that the proportion of individuals showing increased intestinal permeability grows linearly – or quasi – as a function of chronological age, as it was previously reported in *Drosophila melanogaster*. Finally, we validated that, similar to what has been shown in *D. melanogaster*, the Smurf phenotype is a strong indicator of physiological age since it is a harbinger of natural death occurring during normal aging. Interestingly, we show here that Smurfs from different species of drosophila maintained in our laboratory conditions tend to have similar life expectancies – approximately 2.5 days – although the lines they come from show significantly different life expectancies at birth (Fig. 2 and ^9^). In addition, we show that nematode Smurfs from different genetic backgrounds show similar life expectancies too. The fact that only the oldest group of zebrafish Smurfs shows the limit of our group based approach to overcome the limitations associated with the long lifespan of individuals of that species. As we do not have any full longevity curve of these individuals, we cannot know whether the lack of mortality observed in the youngest groups can be due to the young age of Smurfs, as previously presented in ^8^ and ^9^.

Although our present study clearly indicates that the age-related intestinal permeability dramatic increase is an important, evolutionarily conserved step occurring during normal aging, the ultimate causes of ‘natural’ death remain to be determined in those different organisms.

The first description of the Smurf phenotype in drosophila^8^ included numerous other so-called age-related phenotypes. We think that identifying which of these phenotypes are evolutionarily conserved across species will be the first step towards defining the conserved sequences of events driving aging. For example, on the contrary to what was described in *Drosophila melanogaster,* we failed to observe any loss of triglycerides content in nematode where zebrafish Smurfs tend to weigh significantly less than their age-matched non-Smurf counterparts (data not shown). A next step towards identifying events driving age-related death will therefore be to search for the molecular and physiological modifications that are associated with phase 2 of aging we recently described in ^9^. If phase 2 of aging is as broadly present in living organisms as our present study suggests, highly stereotyped molecularly and physiologically^8^ as well as sufficient to explain longevity curves^9^, then we think that identifying the very events responsible for entering into this phase or those characterizing the high risk of impending death associated with that phase could answer fundamental questions about aging and lead to treatments able to significantly improve lifespan/healthspan across a broad range of species including *Homo sapiens.*

## Author contributions statements

MR designed the project. ED, HA and MR designed experimental procedures. ED, LM, LR and MR performed the experiments. ED, HA, JSJ, HT and MR interpreted the results and wrote the manuscript.

## Material and methods

**Smurf assay.** Smurf assay in drosophila was executed as previously described in 8. In order to facilitate the detection of Smurfs in nematodes, blue #1 was replaced by 2.5% (w/v) fluorescein. In zebrafish, Smurfs were assayed by placing individuals in a vat containing 2.5% (w/v) blue #1 in water for 30 minutes. They were then rinsed under clear water until no more blue coloration could be found in the eluate. Fishes showing blue an extended coloration of their body were considered as Smurfs. To calculate the SIR, we plotted the average proportion of Smurfs per vial as a function of chronological age and defined the SIR as the slope of the calculated regression line.

**Animals stocks.** *Drosophila virilis* and *Drosophila mojavensis* were generously provided by Dr Virginie Orgogozo, the different nematode lines were described in ^10^ and the groups of AB zebrafish individuals described in this article were generously provided by Dr Edor Kabashi.

**Rearing conditions.** Flies were cultured in a humidified, temperature-controlled incubator with a 12-h on/offlight cycle at 26 °C in vials containing standard cornmeal medium (0.7% agar, 5.2% Springaline^®^ yeast, 4.3% sucrose, and 2.9% corn flour; all concentrations given in wt/vol). Adult animals were collected under light CO_2_-induced anesthesia, housed at a density of 27–32 flies per vial, and flipped to fresh vials and scored for death every 2–3 d throughout adult life. Smurfs were scored for death daily in nematodes and drosophila.

**Statistical analysis.** Longevity curves as well as Smurf Increase Rates (SIR) were analyzed using GraphPad Prism 6.0. The SIR was calculated by plotting the average proportion of Smurfs per vial as a function of chronological age, with the SIR defined as the slope of the calculated regression line. Each SIR was tested for deviation from both linearity and zero. Binomial test was performed under R 3.2.0 using the function binom.test(x, n, p) with the following parameters x = 4, n = 7 and p = 0/9.

